# Formaldehyde Exposure Induces Systemic Epigenetic Alterations in Histone Methylation and Acetylation

**DOI:** 10.1101/2025.02.25.640236

**Authors:** Jiahao Feng, Chih-Wei Liu, Jingya Peng, Yun-Chung Hsiao, Danqi Chen, Chunyuan Jin, Kun Lu

## Abstract

Formaldehyde (FA) is a pervasive environmental organic pollutant and a Group 1 human carcinogen. While FA has been implicated in various cancers, its genotoxic effects, including DNA damage and DNA-protein crosslinking, have proven insufficient to fully explain its role in carcinogenesis, suggesting the involvement of epigenetic mechanisms. Histone post-translational modifications (PTMs) on H3 and H4, critical for regulating gene expression, may contribute to FA-induced pathogenesis as lysine and arginine residues serve as targets for FA-protein adduct formation. This study aimed to elucidate the effects of FA on histone methylation and acetylation patterns. Human bronchial epithelial cells (BEAS-2B) were exposed to low-dose (100 μM) and high-dose (500 μM) FA for 1 hour, and their histone extracts were analyzed using high-resolution liquid chromatography-tandem mass spectrometry-based proteomics, followed by PTM-combined peptide analysis and single PTM site/type comparisons. We identified 40 peptides on histone H3 and 16 on histone H4 bearing epigenetic marks. Our findings revealed that FA exposure induced systemic alterations in H3 and H4 methylation and acetylation, including hypomethylation of H3K4 and H3K79 and changes in H3K9, H3K14, H3K18, H3K23, H3K27, H3K36, H3K37, and H3R40, as well as modifications in H4K5, H4K8, H4K12, and H4K16. These FA-induced histone modifications exhibited strong parallels with epigenetic alterations observed in cancers, leukemia, and Alzheimer’s disease. This study provides novel evidence of FA epigenetic toxicity, offering new insights into the potential mechanisms underlying FA-driven pathogenesis.

## 1. Introduction

Formaldehyde (FA), one of the simplest but reactive aldehydes, is omnipresent in the environment, leading to significant and often inescapable human exposure. Its pervasive presence stems from both biogenic sources, such as metabolism in plants, organisms, and anthropogenic sources emitted from industrial and residential activities. FA is integral to various industrial applications, which include its incorporation into not only the manufacturing of resin and industrial compounds, including textiles, leather, rubber, cement, and plastic but also into construction, medical, chemical, and pharmaceutical industries. In residential settings, FA is emitted from materials for cabinetry, furniture, and house construction, especially pressed-wood products with FA-related resins, that can contribute continuously to indoor formaldehyde levels and accumulate higher concentrations in poorly ventilated environments, thus posing increased exposure risks. FA is also generated as a by-product of combustion with prominent environmental sources of FA, including tobacco smoke, emissions from motor vehicles, and combustion activities like wood burning. Consequently, due to its diverse applications in human life, human exposure to FA has been widespread in parts of the world. Concentrations of FA indoors were recorded in a range from 12 μg/m^3^ to 23800 μg/m^3^ in different settings including industry, commercial and residential buildings, medical laboratories, and hospitals (Mondal et al., 2024). Previous research reported occupational exposure to FA that industry workers were exposed to FA at a range from 0.18 ppm to 2.37 ppm in a wood processing industry (Wang et al., 2015) and from 0.51 ppm to 2.60 ppm in a utensil factory (Zhang et al., 2010), and laboratory workers at a range from 4.9 μg/m^3^ to 268.7 μg/m^3^ (Pala et al., 2008). Consequently, human exposure to formaldehyde persists as widespread and inadequately addressed public health risk.

Exposure primarily occurs through the inhalation of FA, owing to its ubiquitous presence in the air (Protano et al., 2021). The International Agency for Research on Cancer (IARC) has classified FA as a Group 1 human carcinogen (IARC, 2012). Substantial evidence associates FA exposure with an increased risk of nasopharyngeal cancer in humans (Hauptmann et al., 2004; Marsh et al., 2007; Vaughan et al., 2000) and nasal carcinoma in animals (Kerns et al., 1983; Swenberg et al., 1980). In addition, the toxicological effects of FA exposure related to other diseases have been increasingly documented in both experimental research and epidemiological studies. FA exposure has been reported to induce lung inflammatory disorders (Bhat et al., 2024), liver damages (Othmen et al., 2024), gastrointestinal cancer (TAKAHASHI et al., 1986), cardiovascular diseases and heart development (Zhang et al., 2023), cognitive and neurodegenerative diseases (Li et al., 2016; Mei et al., 2015; Tong et al., 2016), and leukemia (Goldstein, 2011).

However, the molecular mechanisms underlying FA-induced carcinogenesis remain inadequately understood. The accumulation of DNA damage due to FA, with particular emphasis on the formation of DNA adducts and DNA-protein crosslinks (DPCs), has been the focus of elucidating FA carcinogenesis. DPCs were detected in the respiratory tract of rhesus monkeys exposed to FA, corresponding to the sites for nasopharyngeal and nasal tumorigenesis observed in humans (Casanova et al., 1991). DNA cross-links were found to be correlated with FA concentrations and tumor incidence at nasal cavities in FA-exposed rats (Liteplo & Meek, 2003). FA exposure also induced N2-hydroxymethyl-dG monoadducts and dG-dG cross-links in DNA from rat respiratory nasal mucosa (Lu et al., 2010). However, further investigations into the formation of endogenous versus exogenous FA-induced DNA adducts have revealed that the extent of DNA adducts resulting from exogenous FA exposure is only modestly elevated compared to the adducts formed endogenously (Lu et al., 2010, 2011; Moeller et al., 2011; Swenberg et al., 2011). These findings have prompted critical reconsiderations of the role of genetic damage in FA-induced carcinogenesis and indicate that epigenetic mechanisms may contribute substantially to FA-mediated carcinogenicity.

Histone post-translational modification is a major epigenetic mechanism that could be affected by FA exposure. Histones are critical to providing structural support for chromosomes, modulating genomic architecture. Histone modifications, particularly lysine and arginine methylation and lysine acetylation, play crucial roles in regulating transcription, DNA recombination, replication, and repair, maintaining genomic integrity, and contributing to epigenetic memory (Ng et al., 2009). The aberrant histone modifications have been linked to the pathogenesis of various human diseases including cancer (Millán-Zambrano et al., 2022). As an electrophilic compound, FA can form protein adducts by reacting with the nucleophilic side chains of amino acids such as lysine and arginine (Chen et al., 2017; Trézl et al., 2003). Notably, histones are lysine-rich proteins, providing a fertile source for adduction by exogenous FA. FA was found to react with lysine residues to generate the intermediate N^6^-hydroxymethyl-lysine, from which an unstable Schiff base or a stable end product N^6^-formyllysine can be produced (Chen et al., 2017). Interestingly, both Schiff bases and N^6^−formyllysine are resistant to physiological modifications, with N^6^−formyllysine residues found to be refractory to acetylation in cells and in vitro (Edrissi et al., 2013), and FA-induced Schiff bases on lysine residues on histone H4 peptide refractory to acetylation in vitro (Lu et al., 2008). Moreover, recent research demonstrated that chronic elevations in cytosolic FA resulted in significant decreases in H3K4 and HK79 methylations as downstream effects of FA-induced S-adenosylmethionine deficiency (Pham et al., 2023), suggesting that histone methylation is likely to be affected by FA exposure. However, it remains elusive if FA exposure induces systematic changes in methylation and acetylation levels on H3 and H4, increasing the potential for carcinogenesis.

Herein, this exploratory research aspired to decipher and map histone modification patterns induced by FA exposure in vitro, with an emphasis on methylation and acetylation on histones H3 and H4. Bottom-up proteomic approaches, coupled with high-resolution liquid chromatography-tandem mass spectrometry, were applied to detect and accurately measure multiple peptide sequences containing specific modification sites. Compared to immunoblotting methods measuring specific modification sites, this research is capable of revealing modification changes in peptide sequences with multiple modification sites, which advanced the understanding of FA-induced histone modification at the peptide level. Moreover, this research enables comparisons of PTM sites and types induced by different levels of FA exposure, facilitating the exploration of how multiple FA-induced modifications on peptides interact and contribute to changes in histone modification, and potentially on chromatin structure and gene regulation. Ultimately, this research sheds light on understanding the molecular mechanisms contributing to FA carcinogenicity through which FA exposure alters epigenetic signatures on histone H3 and H4, highlighting the potential of histone modifications as biomarkers for FA-induced carcinogenesis.

## 2. Methods

### 2.1. Cell culture and treatment

The immortalized human bronchial epithelial cell line BEAS-2B (CRL-3588) was cultured in DMEM/F-12 (Dulbecco’s Modified Eagle Medium/Nutrient Mixture F-12) (Gibco™, Massachusetts USA), supplemented with 10% fetal bovine serum (FBS) (Corning, NY USA), 100 U/mL penicillin, and 100 µg/mL streptomycin (Gibco™, Massachusetts USA). Cells were maintained under standard culture conditions at 37°C in a humidified incubator with 5% CO₂. Prior to formaldehyde (FA) exposure, cells were seeded at a density of 2 × 10⁶ cells/mL into sterile 75 cm² cell culture flasks and cultured until they reached approximately 85% confluence. The adherent cells were then washed with pre-warmed, sterile phosphate-buffered saline (PBS), and any residual solution was thoroughly removed by vacuum aspiration to prevent dilution of the FA treatments. Fresh FA working solutions were prepared in sterile PBS to achieve target concentrations of 100 µM and 500 µM, representing the low-dose and high-dose exposure groups, respectively. The washed BEAS-2B cells were subsequently exposed to these FA solutions for 1 hour at 37°C. A control group was concurrently included, with cells exposed to sterile PBS without FA to serve as baseline. To ensure the robustness and reliability of results, each experimental condition, including the control, was conducted in six replicates.

### 2.2. Trypan Blue Staining and Cell Viability Assay

Before and after exposure, cells were detached with trypsin from the culture flask surface, centrifuged, and resuspended in PBS solution. An aliquot of cell suspension was mixed thoroughly with 0.4% trypan blue solution at a 1:1 ratio. Stained cell suspension was extracted and loaded on slides into the TC20 automated cell counter. The total number of viable cells was determined by trypan blue staining. Cell viability was determined by calculating the percentage of live cells in five randomly selected microscopic fields. The measurements were repeated three times in each group. The percentage of viable cells was calculated using the formula: % viability = (number of viable cells / total cells counted) × 100. This approach allowed us to determine cell viability with high accuracy for subsequent experimental analyses.

### 2.3. Nuclei Histone isolation and acidic extraction

The overall analytical scheme was summarized in Figure 1A. Following formaldehyde (FA) treatment, cell pellets were subjected to thorough washing with ice-cold phosphate-buffered saline (PBS) (Fisher BioReagents™, USA) to meticulously remove FA-containing PBS and any non-viable cells. The viable cells were subsequently collected by gentle centrifugation. Nuclear isolation lysates (NIL) were meticulously prepared using a Tris-HCl buffer system comprising 15 mM Tris, 60 mM KCl, 15 mM NaCl, 5 mM MgCl₂, 1 mM CaCl₂, 250 mM sucrose, 1 mM dithiothreitol (DTT), 5 nM microcystin, 10 mM sodium butyrate, 500 μM AEBSF, and 0.2% NP-40. The washed, viable cells were promptly resuspended in NIL buffer and incubated on ice for 10 minutes to facilitate nuclear lysis. Following centrifugation, supernatants were carefully discarded, and the resulting cell pellets, enriched with chromatin-bound histones, were collected in vials.

**Figure 1.**
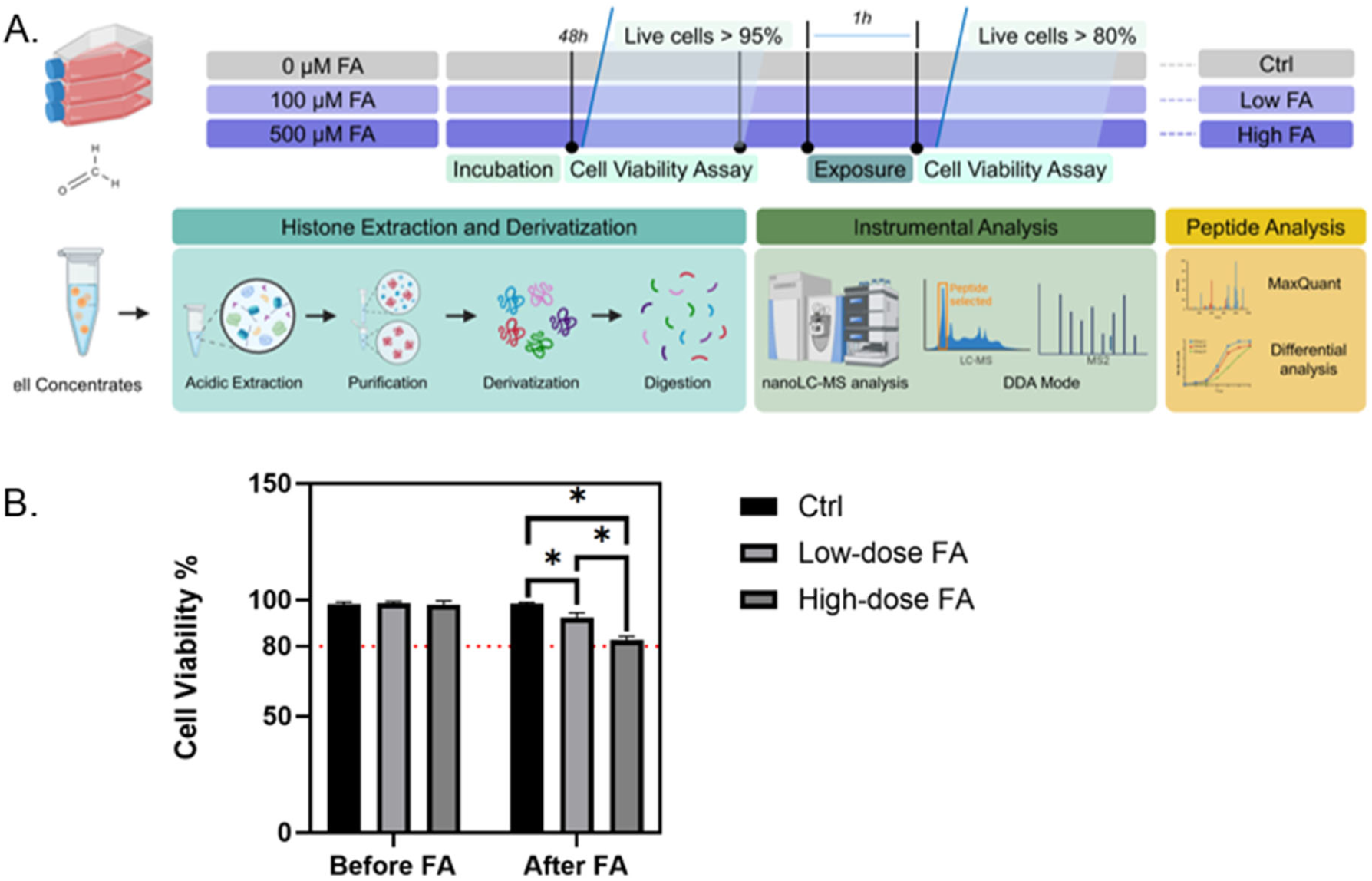
A. Experimental workflow started with FA treatment, sample harvest, and histone extraction of cells among three groups: control, low-dose, and high-dose (*N* = 6). B. Cell viability test before FA exposure and after FA exposure. The cell viability was above 80% in three groups after 1h FA exposure. (*p < 0.05).

Histones were isolated from nuclei by treating the nuclear pellet with 0.4 N sulfuric acid, which was carefully introduced to facilitate the selective extraction of histones. The mixture was then incubated at 4°C for 2 hours to maximize extraction efficiency. Following incubation, the sample underwent centrifugation, and the supernatant was subsequently treated with trichloroacetic acid, thereby inducing histone precipitation. The resulting histone precipitate was subjected to a thorough purification process, including multiple washes with acetone to effectively remove contaminating impurities. The purified histone samples were then meticulously collected and stored for downstream analyses.

### 2.4. Histone derivatization and enzymatic digestion

Histone samples were prepared for derivatization by aliquoting 10 µg of each sample into an ammonium bicarbonate buffer (pH 8.0). A fresh propionylation reagent, prepared by mixing propionic anhydride with acetonitrile in a 1:3 (v/v) ratio, was carefully introduced into each histone sample in a 1:4 (v/v) ratio. The derivatization reaction proceeded for 15 minutes at room temperature, after which the samples were dried and reconstituted. The derivatization procedure was repeated to ensure a propionylation completion rate of greater than 95%.

After verifying that the pH remained at 8.0, reconstituted histone samples were subjected to proteolytic digestion with trypsin, followed by incubation at 37°C for 17 hours to achieve complete enzymatic cleavage. The digestion reaction was halted by freezing the samples at −80°C. Subsequent desiccation and reconstitution in ammonium bicarbonate buffer (pH 8.0) were performed, followed by a secondary propionylation step to modify the N-termini of the histone peptides. Samples were then dried and desalted to remove impurities. Peptide concentrations in each desalted sample were then rigorously quantified using a NanoDrop spectrophotometer, ensuring consistency and accuracy across samples before advancing to downstream analyses.

### 2.5. LC-MS/MS analysis

The LC-MS/MS analysis was performed using an UltiMate 3000 RSLCnano system coupled to a Q Exactive HF Orbitrap mass spectrometer through an EASY-Spray ion source (ThermoFisher Scientific). A binary solvent system (solvent A: 0.1% FA in water; solvent B: 0.1% FA in acetonitrile) was used for peptide separation at a flow rate of 250 nL/min. Peptides (0.3μg/μL, 6 μL) were separated on a PepMap C18 analytical column (2 μm particle, 50 cm×75 μm i.d., ThermoFisher Scientific). LC separation was performed using the following gradient: held at 4% B for 4 min, from 4% to 8% B in 0.1 min, 8% to 40% B in 93 min, from 40% to 95% in 0.1 min, held at 95% B for 10 min, 95% to 4% B in 0.1 min, and held at 2% B for 17 min for re-equilibrating column. MS and MS/MS spectra were acquired in profile mode using a data-dependent top-15 method and the resolution for a full MS scan (350 to 1,650 m/z) was set to 60,000 with a maximum fill time of 50 ms. Precursors were isolated with a window of 1.4 m/z and fragmented with higher-energy collisional dissociation (HCD, normalized collision energy of 27). Resolution for MS/MS spectrum was set to 30,000 with a maximum full time of 50 ms. The AGC (automatic gain control) targets for MS and MS/MS scans were 1×10^6^ and 2×10^5^, respectively. Precursor ions with single, seven, and higher charge states were excluded from fragmentation, and dynamic exclusion time was set to 20 s.

### 2.6. Data analysis

Data was loaded on Maxquant (Version 2.2.0.0.) for the best spectrum identification on each unique peptide containing the specific post-translational modification. In the group-specific parameters, the variable modifications contained unmodified lysine; lysine mono-methylation, lysine di-methylation, and lysine tri-methylation. The N-terminal of histone propionylation was set as a fixed modification. In the configuration, the parameters were modified to match the peptide with an additional propionyl group. Due to histone propionylation, the patterns of trypsin digestion will mimic the digestion pattern of the Arg-C enzyme. Thus, the digestion mode was set as specific Arg-C digestion. The database was imported from the UniProt protein database containing complete protein coverage of homo sapiens (UP000005640). The minimal peptide length was set as 5 amino acids, and the maximal peptide mass was set as 4600 Da. Regarding the accuracy of peptide identification, the false detection rate (FDR) at the peptide spectrum match level, which is determined by the target-decoy approach, and at protein level, was set as 0.01, respectively.

With the identified best MS/MS spectrum for each unique peptide, the retention time and mass/charge ratio of typically fragmented ions present in the MS/MS spectrum were applied to retrieve the peptides in full scan, followed by quantifying the signal area in the full scan detection. The log-transformed signal area was collected for peptide analysis. The differences among the three groups were analyzed using non-parametric one-way ANOVA, as well as multiple student t-tests to statistically compare the differences among specific peptides.

## 3. Results

### 3.1. Peptides identification on LC-MS/MS

Peptide identification was primarily facilitated through tandem mass spectrometry (MS/MS) spectrum, fragmentation of digested histone proteins produces a spectrum of smaller ions, specifically b-ions and y-ions, which represent fragments generated from the N-terminal and C-terminal regions of the peptide, respectively. The MS/MS spectrum was analyzed to uncover all ions in a peptide sequence, providing a detailed mass-to-charge (m/z) ratio that correlates to specific amino acid residues within the peptide sequence. The identification process involves matching the observed fragment ions on MS/MS spectra against protein sequences to generate known peptide sequences on histones.

For instance, the peptide _73_EIAQDF**K_2me_**TDLR_83_, which includes a di-methylation modification, was identified through the analysis of multiple b-ions and y-ions derived from the peptide fragmentation. Each of these ions corresponds to a specific segment of the peptide, and together, they allow for the precise determination of the sequence, as well as the identification of post-translational modifications (PTMs) (Figure 2A). The unique patterns of fragmentation observed in the MS/MS spectrum provide critical information that confirms the presence of specific amino acids and the location of modifications.

**Figure 2A.**
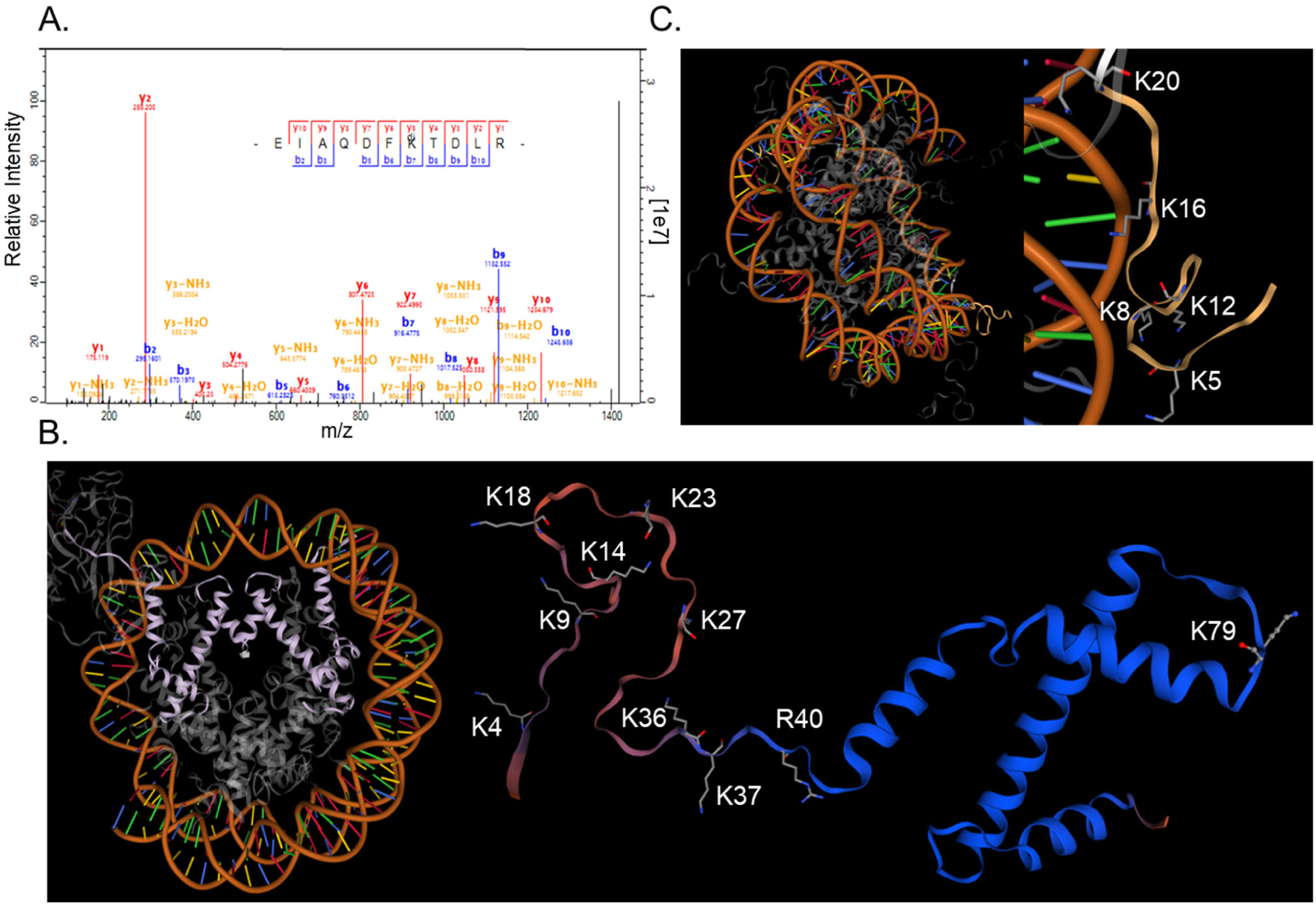
Representative peptide identification procedure using H3 _73_EIAQDF**K_2me_**TDLR_83_ as an example. MS/MS spectrum to identify the peptide sequence with a match between b-ions and y-ions; B. Modification sites of Histone H3 on lysine and arginine that are associated with FA exposure with statistical significance (p<0.05), identified by intensities of PTM-combined peptides. C. Modification sites of Histone H4 on lysine and arginine that are associated with FA exposure with statistical significance (p<0.05), identified by intensities of PTM-combined peptides.

Histone H3 (UniProt ID P84243) is a primary target for extensive post-translational modifications (PTMs), including methylation and acetylation on lysine and methylation on arginine residues (Figure 2B). Using high-resolution LC-MS/MS analysis, a total of 40 peptide fragments exhibiting specific PTMs were detected (Table 1). Among these, lysine modifications were observed on 39 peptides: 4 peptides contained demethylation modifications on lysine residues, 19 peptides had mono-methylation, 12 peptides exhibited di-methylation, 8 peptides had tri-methylation, and 19 peptides showed lysine acetylation. Additionally, arginine methylation was detected in 4 peptides, with two peptides showing arginine mono-methylation and two peptides showing arginine di-methylation.

**Table 1.**
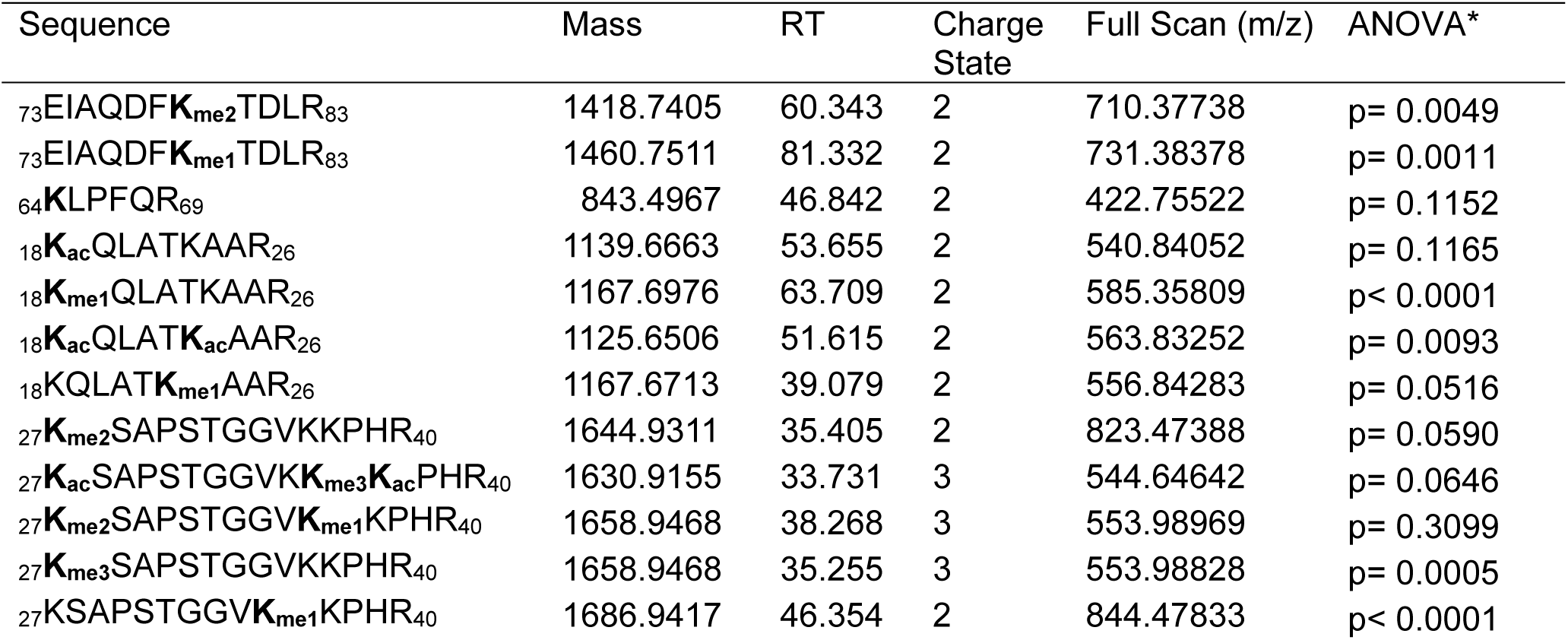
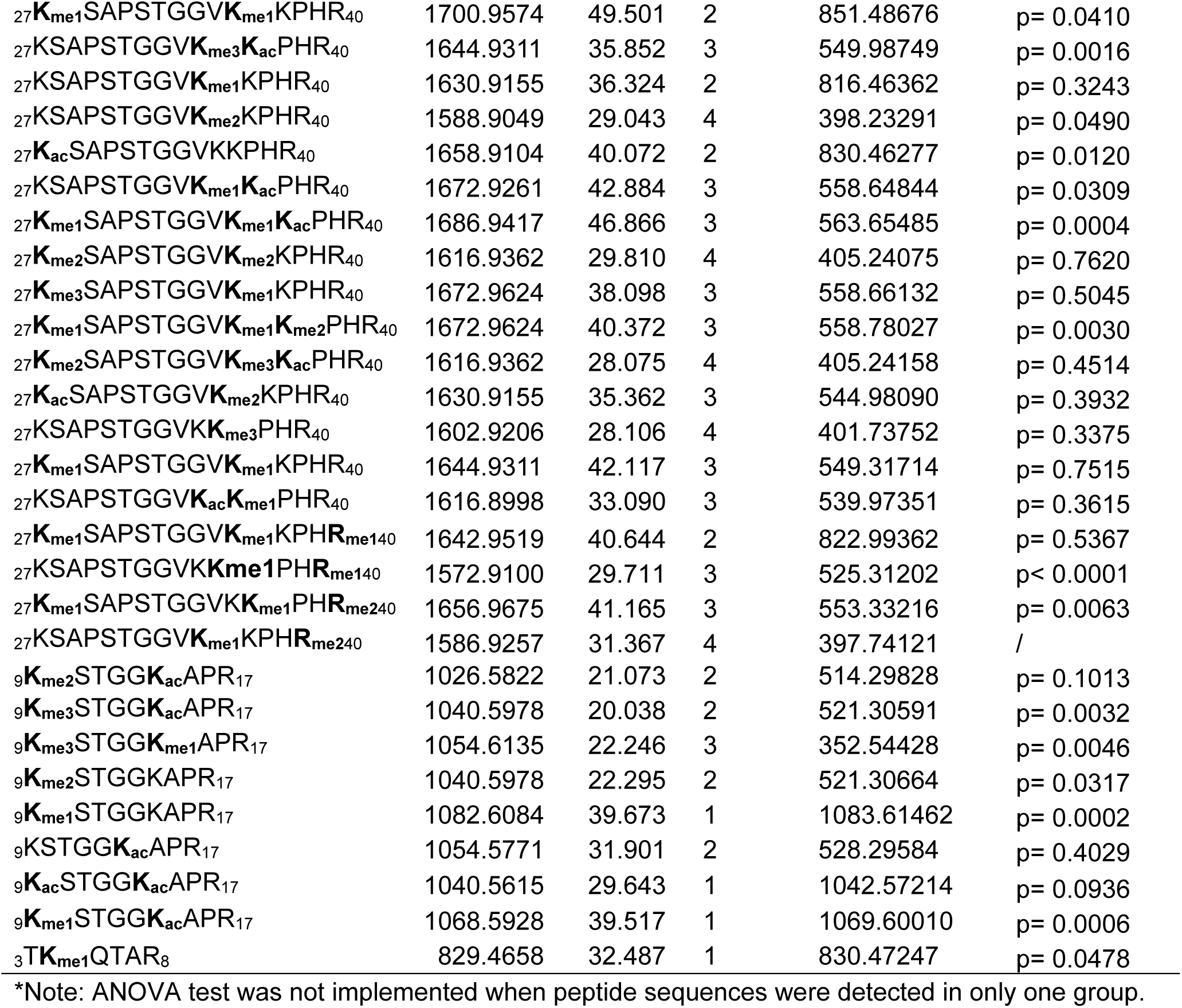
PTM Peptide Fragments on Histone 3 Observed by LC-MS/MS.

On histone H4 (UniProt ID: P62805), a total of 16 peptide sequences were identified (Figure 2C; Table 2). Among these peptides, 13 contained lysine acetylation (K_ac_), 8 exhibited mono-methylation (K_me1_), and 1 contained di-methylation (K_me2_). Interestingly, no arginine methylation was detected in any of the identified peptides. The peptide sequences studied were found to include both single and multiple modifications at various lysine residues.

**Table 2.** PTM Peptide Fragments on Histone 4 Observed by LC-MS/MS.

| Sequence | Mass | RT | Charge State | Full Scan (m/z) | ANOVA* |
| --- | --- | --- | --- | --- | --- |
| <sup>4</sup>GK<sub>ac</sub>GGK<sub>ac</sub>GLGK<sub>ac</sub>GGAK<sub>ac</sub>R<sub>17</sub> | 1493.8314 | 42.023 | 2 | 747.92322 | p= 0.2667 |
| <sup>4</sup>GKGGKGLGK<sub>me1</sub>GGAK<sub>ac</sub>R<sub>17</sub> | 1493.8678 | 41.026 | 3 | 498.96313 | p= 0.2001 |
| <sup>4</sup>GK<sub>ac</sub>GGK<sub>ac</sub>GLGK<sub>ac</sub>GGAKR<sub>17</sub> | 1507.8471 | 44.335 | 2 | 754.93121 | p= 0.5377 |
| <sup>4</sup>GKGGKGLGK<sub>ac</sub>GGAK<sub>ac</sub>R<sub>17</sub> | 1521.8627 | 47.637 | 2 | 761.93842 | p= 0.0003 |
| <sup>4</sup>GKGGKGLGK*GGAK<sub>ac</sub>R<sub>17</sub> | 1535.8784 | 50.372 | 1 | 1536.88647 | p< 0.0001 |
| <sup>4</sup>GK<sub>me1</sub>GGK<sub>ac</sub>GLGKGGAKR<sub>17</sub> | 1549.8940 | 52.783 | 1 | 1550.90173 | p< 0.0001 |
| <sup>4</sup>GK<sub>ac</sub>GGK<sub>me1</sub>GLGK<sub>ac</sub>GGAK<sub>ac</sub>R<sub>17</sub> | 1521.8627 | 47.193 | 2 | 761.93835 | p< 0.0001 |
| <sup>4</sup>GK<sub>me1</sub>GGK<sub>ac</sub>GLGKGGAK<sub>ac</sub>R<sub>17</sub> | 1535.8784 | 51.446 | 1 | 1536.88440 | p< 0.0001 |
| <sup>4</sup>GKGGKGLGK<sub>ac</sub>GGAK<sub>ac</sub>RHR<sub>19</sub> | 1758.9965 | 41.090 | 2 | 881.00879 | p= 0.3062 |
| <sup>1</sup>SGRGKGGKGLGK<sub>ac</sub>GGAK<sub>ac</sub>R<sub>17</sub> | 1822.0173 | 39.257 | 2 | 912.01654 | p= 0.1657 |
| <sup>1</sup>SGRGKGGKGLGKGGAK<sub>ac</sub>R<sub>17</sub> | 1836.0330 | 42.596 | 2 | 919.02521 | p< 0.0001 |
| <sup>1</sup>SGRGK<sub>me2</sub>GGKGLGKGGAKR<sub>17</sub> | 1822.0537 | 39.867 | 2 | 912.03314 | / |
| <sup>1</sup>SGRGK<sub>me1</sub>GGK<sub>ac</sub>GLGKGGAK<sub>ac</sub>R<sub>17</sub> | 1836.0330 | 44.620 | 2 | 613.01959 | p< 0.0001 |
| <sup>1</sup>SGRGK<sub>me1</sub>GGKGLGKGGAK<sub>ac</sub>R<sub>17</sub> | 1850.0486 | 52.246 | 2 | 926.02716 | p= 0.5330 |
| <sup>1</sup>SGRGK<sub>me1</sub>GGKGLGKGGAKR<sub>17</sub> | 1864.0643 | 44.201 | 3 | 622.36298 | p= 0.0211 |
| <sup>20</sup>K<sub>me1</sub>VLRDNIQGITKPAIR<sub>35</sub> | 2003.1891 | 71.115 | 3 | 668.73773 | p= 0.0520 |

\*Note: ANOVA test was not implemented when peptide sequences were detected in only one group.

### 3.2. Formaldehyde induces H3K4 and H3K79 hypomethylation

Lysine K4 is the first lysine residue from the N-terminal that can undergo covalent modification on histone H3. FA exposure induces targeted alterations in methylation states at lysine residues K4 on histone H3, as evidenced by peptides _3_TKQTAR_8_ (Table 1). Exposure to FA led to a statistically significant reduction of H3K4me1 by 2.522% (p<0.01) and 1.411% (P<0.05) by low- and higher dose of FA, respectively, compared to normal states, implying a slight inhibition of H3K4me1 by FA (Figure 3A).

**Figure 3.**
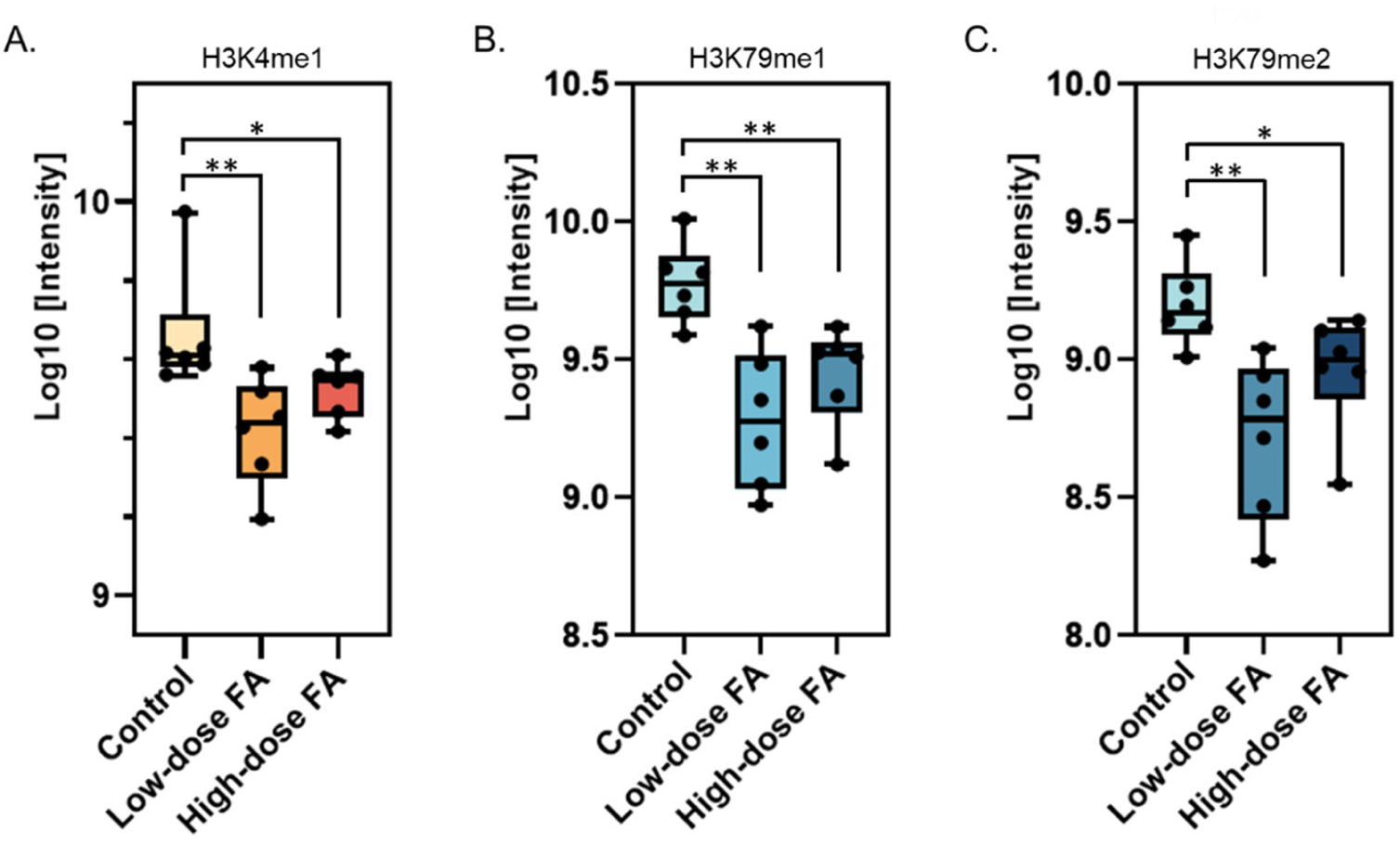
Peptide Intensity analysis on methylation states of H3K4me1 (A), H3K79me1(B), and H3K79me2 (C) in control, low-dose and high-dose groups (N = 6). (\**p* < 0.05, \*\**p* < 0.01).

H3K79 is uniquely located within a loop in the globular domain exposed on the nucleosome surface, occupying a position in the ordered core domain of histone H3 within a short turn connecting the first and second helices of the conserved histone fold (Luger et al., 1997; Min et al., 2003). FA exposure induces changes in both mono-methylation (H3K79me1) and di-methylation (H3K79me2) levels, assessed through the intensity of peptide sequences _73_EIAQDFKTDLR_83_ (Table 1). Similar to H3K4me1, H3K79 mono- and di-methylation were shown statistically significant decreases in low-dose FA exposure to 94.930% and 94.765% (p<0.01) (Figure 3B; 3C), respectively, as compared to the control. The levels of H3K79me and H3K79me2 were lower in the high-dose FA-treated group compared to the control group, albeit no dose-dependent response was observed.

### 3.3. Formaldehyde regulates both methylation and acetylation on H3K9, H3K14, H3K18, and H3K23

Besides H3K4 methylation on the N-terminal of histone H3, FA exposure induced covalent modification on K9 and K14 at the N-terminal of histone H3 (Figure 4A). Acetylation and methylation of H3K9 and H3K14 are key regulators of chromatin structure and gene expression, although H3K14 methylation is less well studied. FA exposure induces significant alterations in peptides that contain modifications on H3K9 and H3K14, revealed by peptide sequences that contained modification marks on either or both lysine, which were identified as H3K9me1K14, H3K9me1K14ac, H3K9me2K14, H3K9me2K14ac, H3K9me3K14me1. Peptides containing H3K9me2K14 and H3K9me3K14ac were the most abundant under control conditions among five H3K9K14 combined peptides (Figure 4A). After low or high FA exposure, these two modifications remained in the highest abundance, even though statistically significant depletion was witnessed. H3K9me3K14me1 was the least abundant peptide sequence, both in the control group and FA-exposed groups. Following exposure to low-dose FA, statistically significant reductions in all five peptides were observed to a level ranging from 95.396% to 96.565% (p<0.05) compared to non-exposed groups, suggesting a modest repressive effect of FA on these modifications. At 500 μM FA exposure, the suppression of modification-contained peptides due to FA was less pronounced than 100 μM FA exposure compared to the reference levels, depicting a slight elevation in intensities at 500 μM FA exposure compared to 100 μM FA exposure (Figure 4A). Among these five peptide species, peptides containing H3K9me2K14 restored their level similar to the reference level, while intensities of the other four peptides were still statistically lower in high-FA exposure than non-exposed levels (p<0.01). These findings concluded that FA exposure led to a reduction in methylation and acetylation on H3K9 and H3K14 to an extent that was inconsistently related to FA concentrations, as suppression was more evident at low-dose exposure rather than high-dose FA exposure.

**Figure 4.**
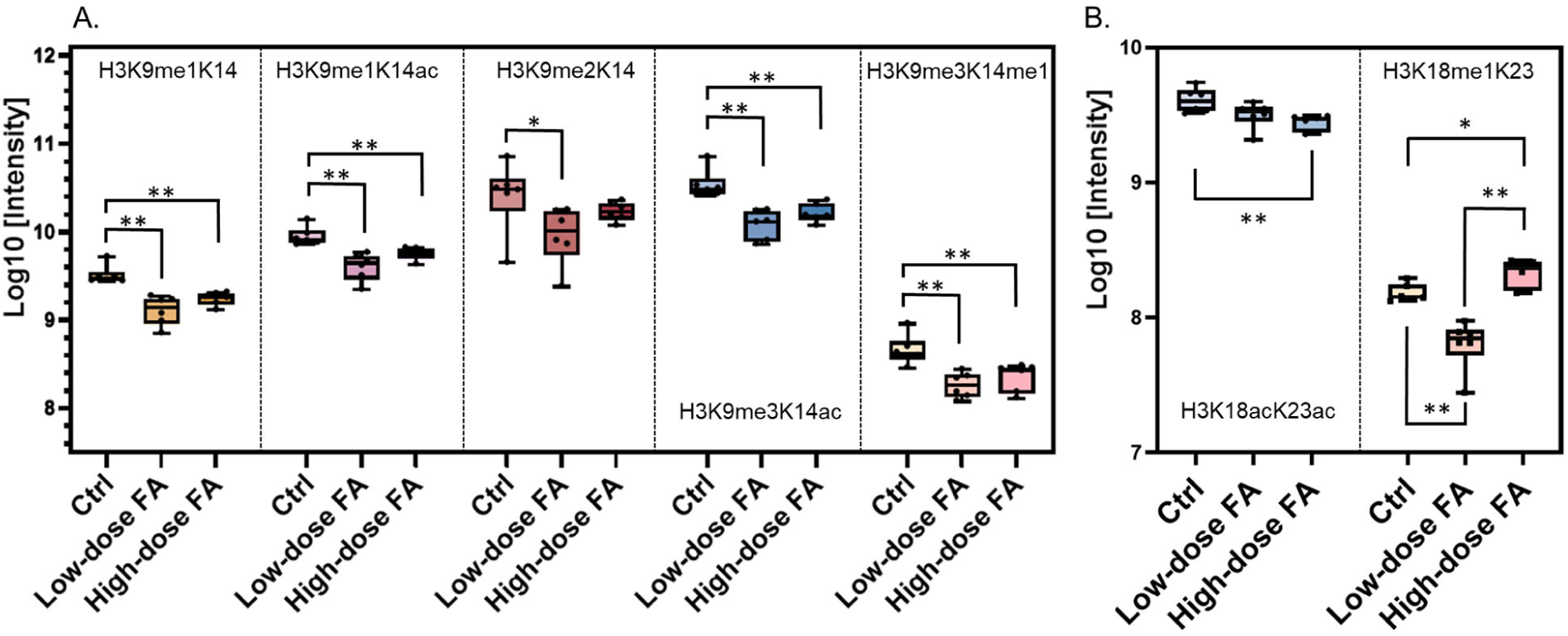
Peptide Intensity analysis on combined PTM states of H3K9K14 (A) and H3K18K23 (B) in control, low-dose and high-dose groups (N = 6). (\**p* < 0.05, \*\**p* < 0.01).

Lysine residues K18 and K23 on histone H3 are crucial sites for transcriptional activation. Modifications on H3K18 and H3K23 were revealed by peptide sequences consisting of H3K18acK23ac and H3K18me1K23 (Figure 4B). The global level of H3K18acK23ac, both in non-FA and FA-treated groups, was significantly higher than H3K18me1K23. Exposure to 100 μM FA resulted in a modest reduction, bringing H3K18me1K23 to 95.346% (p<0.01) and H3K18acK23ac to 98.882%, suggesting that low-dose FA exposure can significantly decrease these two modifications. However, at high-dose FA exposure, these two peptides exhibited different patterns. The intensity of H3K18acK23ac at high FA dose declined further than in the control and low-dose FA groups to 98.259% (p<0.01), but the intensity of H3K18me1K23 was elevated than in the low-dose FA group, even reaching a level (101.729%, p<0.05) slightly higher than the control group. This discrepancy implied that high-concentration FA increased the global level of H3K18me1K23 but slightly decreased peptides containing H3K18acK23ac.

### 3.4. Formaldehyde exposure regulates modifications of H3K27, H3K36, and H3K37

From proteomic analysis, nine peptide sequences containing different epigenetic marks on H3K27, H3K36, and H3K37 were confirmed to be altered by FA exposure (p<0.05) (Table 1). Under unexposed conditions, H3K27K36me3K37ac, H3K27K36me1K37, and H3K27me1K36me1K37ac were the top 3 abundant peptide sequences, while the top 3 peptide abundance in low-dose FA group shifted to H3K27K36me1K37, H3K27K36me3K37ac, and H3K27me1K36me1K37me2. In the high-dose FA group, peptides containing H3K27me1K36me1K37me2, H3K27K36me3K37ac, and H3K27me3K36K37 had the highest intensities (Figure 5).

**Figure 5.**
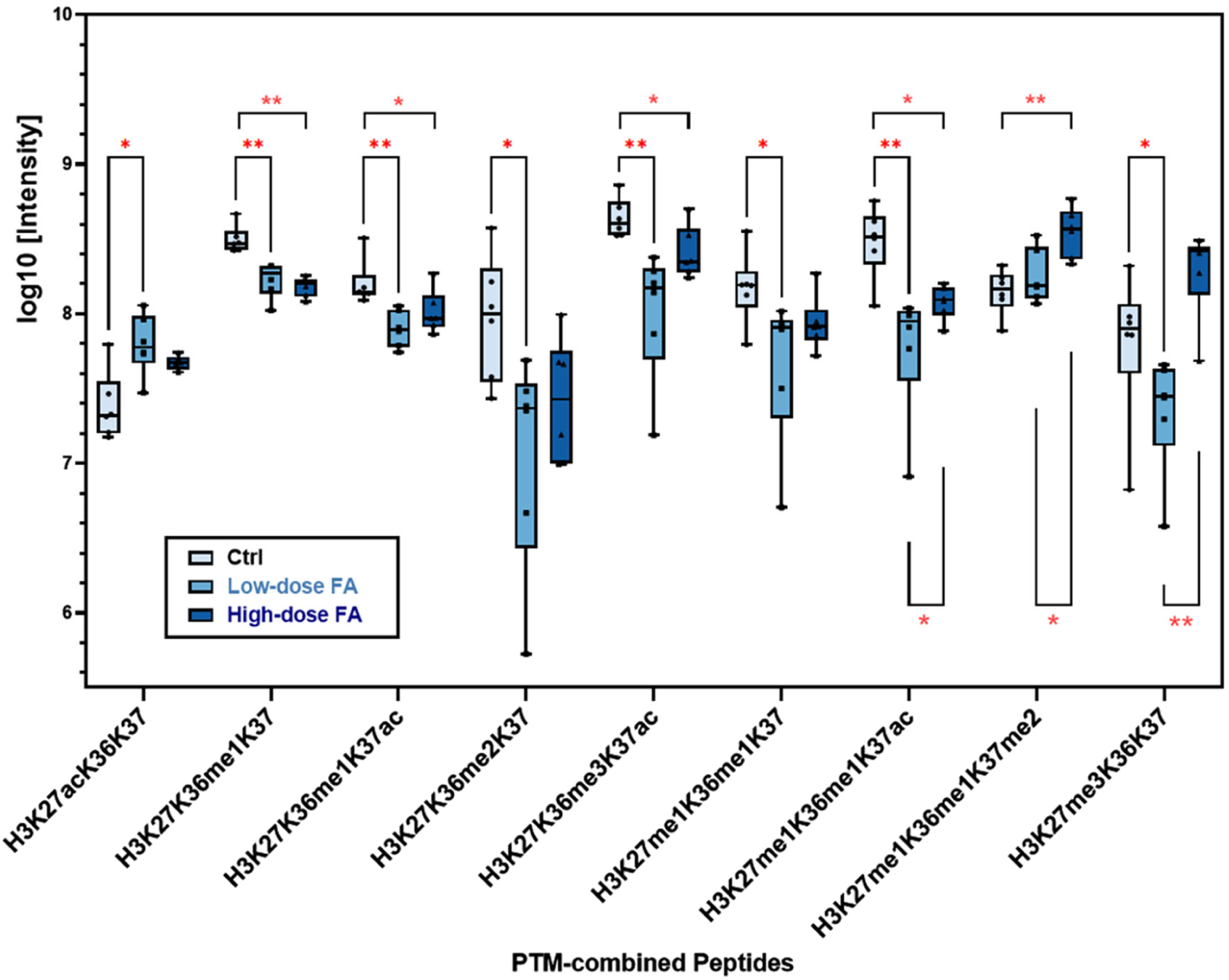
Peptide Intensity analysis on combined PTM states of H3K27, H3K36, and H3K37 in control, low-dose and high-dose groups (N = 6). (\**p* < 0.05, \*\**p* < 0.01).

Histone peptides containing H3K27me1K36K37, H3K27K36me1K37ac, H3K27K36me2K37, H3K27K36me3K37ac, H3K27me1K36me1K37, H3K27me1K36me1K37ac, and H3K27me3K36K37 shared a similar pattern in FA exposure. Compared to the control baseline level, these seven peptides declined in intensities with statistical significance (p<0.05) in low-dose FA groups (Figure 5). Comparing the magnitude of peptide intensities between high-dose group and low-dose group to control level, the peptide sequence containing H3K27K36me1K37 remained unchanged at 96%, while peptides H3K27K36me1K37ac, H3K27K36me2K37, H3K27K36me3K37ac, and H3K27me1K36me1K37 exhibited a slight increase in high-dose FA exposure than in low-dose FA exposure even though statistical significance was not established. Peptide H3K27me1K36me1K37ac exhibited an elevation of intensity in high-dose FA exposure (95.250%) than in low-dose FA exposure (91.643%), while its intensity in high-dose FA exposure is still lower than in control with statistical significance (p<0.05). Peptide H3K27me3K36K37 in high-dose FA exposure groups (106.269%) exhibited a dramatic increase in intensity than in low-dose group (94.150%), even moderately exceeding the control level, although statistical significance was not established in the discrepancy between the control and high-dose FA group.

On the contrary, the peptides with modifications H3K27acK36K37 and H3K27me1K36me1K37me2 were promoted in the FA-treated group than in the control. As the peptide lowest in intensity in control, peptide H3K27acK36K37 increased in the low-dose group (105.641%, p<0.05) and high-dose group (103.937%), suggesting that FA exposure increased acetylation in H3K27. As the only peptide that contained methylation marks on all three lysine residues K27, K36, and K37, peptide H3K27me1K36me1K37me2 showed a slight increase in the low-dose exposure group (101.264%), and a dramatic increase in the high-dose group (104.872%), which statistical significance was reported between high-dose exposure and control (p<0.01), and between high-dose exposure and low-dose exposure (p<0.05).

### 3.5. Formaldehyde modulates methylation on histone H3 at arginine R40

Peptide sequence analysis identified four distinct sequences containing arginine methylation epigenetic modifications. Intriguingly, the presence of arginine methylation was invariably accompanied by mono-methylation at lysine H3K36 across all four peptides. Peptide sequences bearing the H3K27K36me1R40me1 modification exhibited relatively consistent levels across the three experimental groups. However, a progressive diminution in the abundance of H3K27K36me1R40me1 was observed with escalating FA concentrations, indicative of a dose-dependent response of FA exposure on decreasing this peptide. Specifically, the intensity declined from the control group to 93.796% in the low-dose group (p<0.01) and further decreased to 88.634% in the high-dose group (p<0.01) (Figure 6).

**Figure 6.**
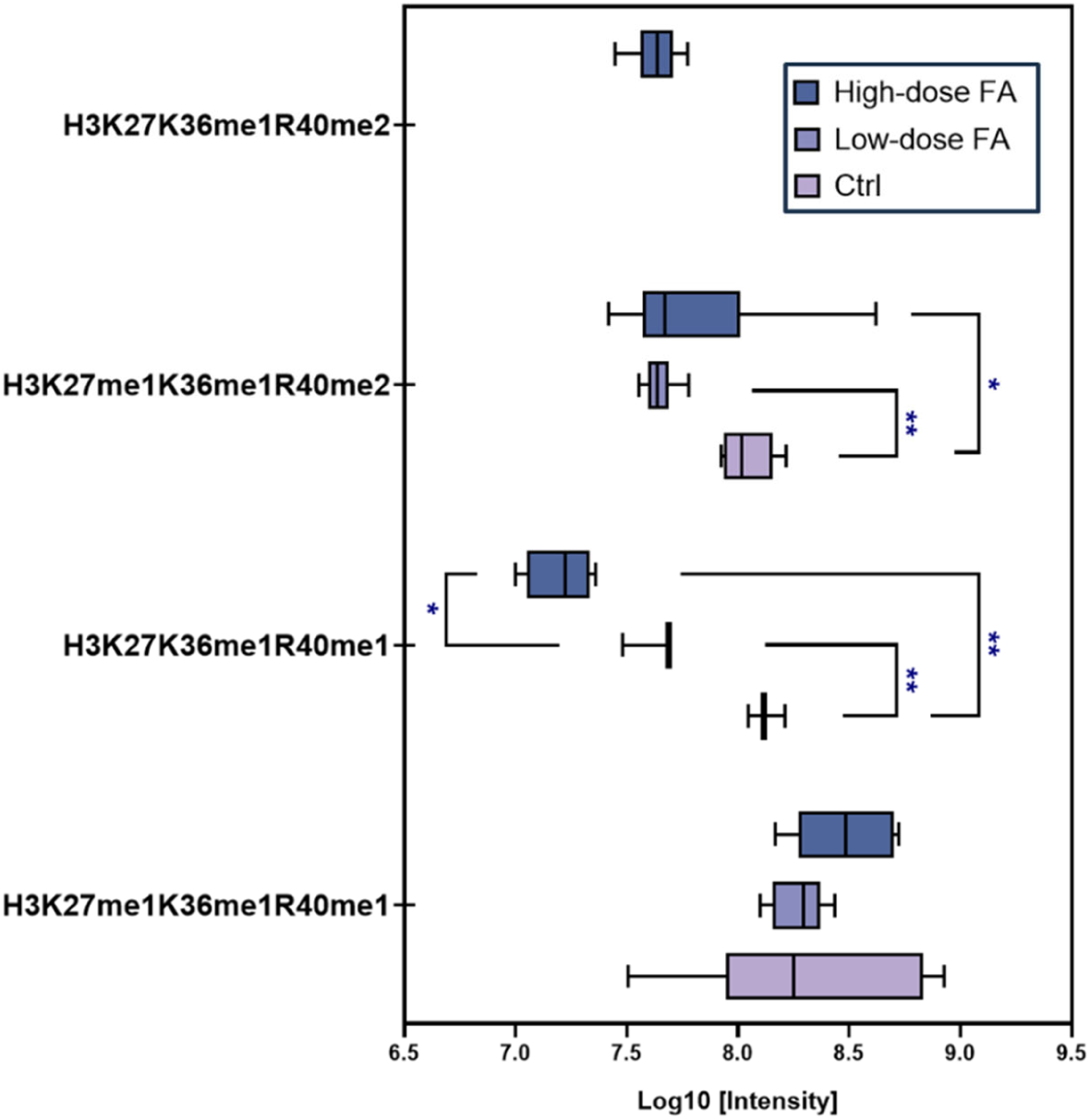
Peptide Intensity analysis on peptides containing PTMs occurring on arginine residues of histone H3 (H3R40) in control, low-dose and high-dose groups (N = 6). (\**p* < 0.05, \*\**p* < 0.01).

In contrast, the peptide containing H3K27K36me1R40me2 emerged exclusively in the high-dose FA exposure group, with its presence absent in both the control and low-dose groups. This observation strongly suggests that higher FA concentrations may catalyze an additional methylation event at arginine R40, thereby driving the conversion of H3R40me1 to H3R40me2. In summary, these findings suggest that FA exposure precipitates a reduction in H3K27K36me1R40me1, while concurrently promoting di-methylation at H3R40, culminating in the formation of H3K27K36me1R40me2. This evidence underscores the potential role of FA in orchestrating site-specific epigenetic alterations through enhanced arginine R40 methylation.

### 3.6. Effects of Formaldehyde on methylation and acetylation of histone H4 at lysine K5, K8, K12, and K16

In this study, we observed notable impacts of FA exposure on the acetylation and methylation landscape of histone H4, specifically targeting lysine residues K5, K8, K12, and K16. Eight peptide sequences containing combined modification states on these four lysine residues were identified as strongly associated with FA exposure. Except for H4K5me1K8K12K16, all peptides declined in intensities in both high-dose FA exposure and low-dose FA exposure compared to control, further evaluated with strong statistical significance (Figure 7).

**Figure 7.**
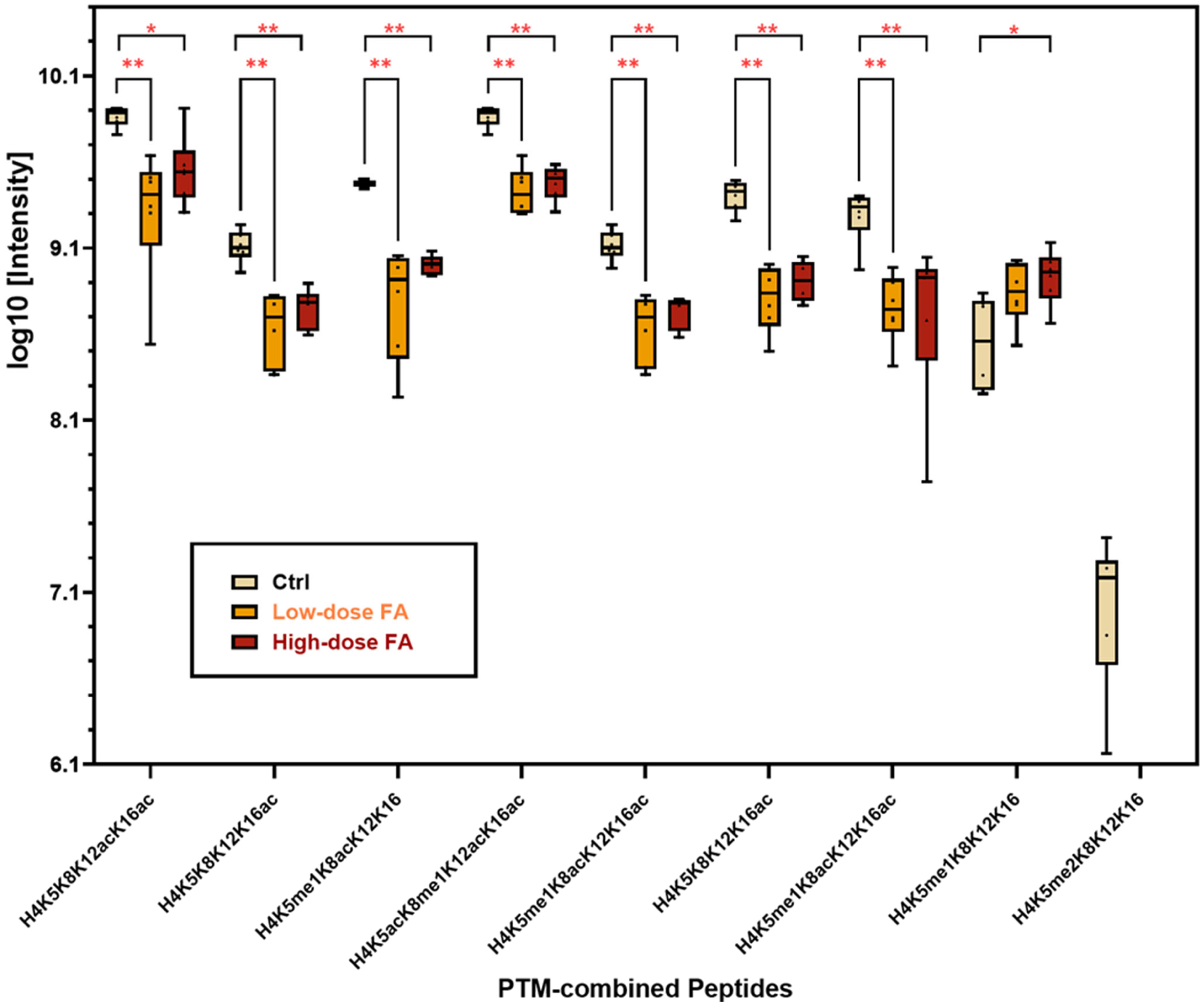
Peptide Intensity analysis on combined PTM states of H4K5, H4K8, H4K12, and H4K16 in control, low-dose and high-dose groups (N = 6). (\**p* < 0.05, \*\**p* < 0.01).

Peptides consisting of H4K5K8K12acK16ac and H4K5K8me1K12acK16ac were the most abundant peptides consisting of epigenetic marks on H4K5, K8, K12, and K16 among three groups. Even though FA exposure lowered the intensity of these two peptides, their intensity remained the highest in FA-treated groups, indicating that combined modifications of H4K5K8K12acK16ac and H4K5K8me1K12acK16ac were the most common epigenetic marks in peptides. Peptides containing H4K5me1K8acK12K16 (92.676%, p<0.01), H4K5K8K12K16ac (93.651%, p<0.01), and H4K5me1K8acK12K16ac (94.159%, p<0.01) showed the greatest decline when exposed to low-dose FA. Besides, peptides containing H4K5K8K12acK16ac, H4K5K8K12K16ac, H4K5K8me1K12acK16ac, and H4K5me1K8acK12K16ac were significantly declined in low-dose groups in a range between 94.314% and 95.596% (p<0.01). Among these seven PTM-combined peptides, high-dose FA exposure did not induce significant changes in the other seven peptides compared to low-dose FA exposure, which indicated that impacts of FA exposure on these peptides did not reveal a dose-dependent pattern, and increasing FA concentrations might not induce further modification on lysine residues of histone H4.

Methylation of H4K5 was affected by FA exposure. H4K5me1K8K12K16 was the only modification that exhibited a slight increase with the elevation of FA, which reached 103.437% in the low-dose group and 104.524% in the high-dose group. In the low-dose FA exposure group, the magnitude of facilitation of this H4K5me1 was not statistically significant. However, this modification in the high-dose group is significantly (p<0.05) higher than in the control group, demonstrating that high-dose FA exposure has promotion effects on H4K5me1. Differently, H4K5me2K8K12K16 was the only PTM-combined peptide presented in the control group but not in any of the FA-exposed groups. These results further suggest that FA exposure eliminated di-methylation of H4K5me2 and increased mono-methylation of H4K5me1.

### 3.7. Effects of FA on single PTM marks on histone H3 and H4

Peptide analysis performed in this study was an integral analysis of peptide sequences that contained multiple PTMs affected by FA exposure, but alterations in single PTMs might further imply the sites and PTM types due to FA interference. Therefore, we explored further to map a single PTM and its sites by leveraging intensities of peptide sequences that exhibited statistical significance being affected by FA exposure. Our results showed ten PTMs on histone H3 were negatively associated with FA exposure, which were H3K4me1, H3K9me1, H3K9me2, H3K9me3, H3K14me1, H3K14ac, H3K36me3, H3K37ac, H3K79me1, and H3K79me2, depicted as lower intensities were quantified in low-dose FA exposure and high-dose FA exposure with statistical significance (p<0.05) compared to the non-exposed baseline level (Figure 8A). These PTMs remained at significantly lower intensities in high-dose FA exposure than in non-exposed conditions (p<0.05). Besides, H3K37ac was the PTM type that exhibited a significant difference between the low-FA and high-FA groups, with its intensity being significantly higher in the high-FA than the low-FA group, indicating a higher concentration of FA contributed less to its reduction. H3K27me1 and H3K36me2 were the PTM marks that significantly decreased in low-FA exposure (95.395% and 88.494%, p<0.05) but reached similar levels as baseline levels in high-FA exposure. Especially, H3K27me1 exhibited a statistical difference between low-FA exposure and high-FA exposure, with its intensity being significantly higher in the high-FA than low-FA group, indicating a higher concentration of FA contributed less to its reduction, similar to H3K37ac. H3K18ac, H3K23ac, and H3R40me1 exhibited a significant reduction in intensities only when exposed to high FA, while H3K36me1 and H3K37me2 exhibited significant elevation in intensities only when exposed to high FA. Differently, when exposed to low or high FA, H3K27ac was consistently elevated, indicating that FA exposure facilitated its acetylation. Histone modifications on H3K18me1, H3K27me3, and H3R40me2 exhibited opposite changes under FA exposure, in which low FA exposure decreased their intensities and high FA exposure increased their intensities, implying that these PTMs were sensitive to FA exposure and their dose-response is not linear. Furthermore, analysis of single PTM marks of histone H4 displayed that H4K5me1, H4K5ac, H4K8ac, H4K8me1, H4K12ac, and H4K16ac were exceptionally inhibited by FA, both in low-dose and high-dose (Figure 8B). However, the suppression due to high-FA exposure is slightly moderate rather than low-FA exposure.

**Figure 8.**
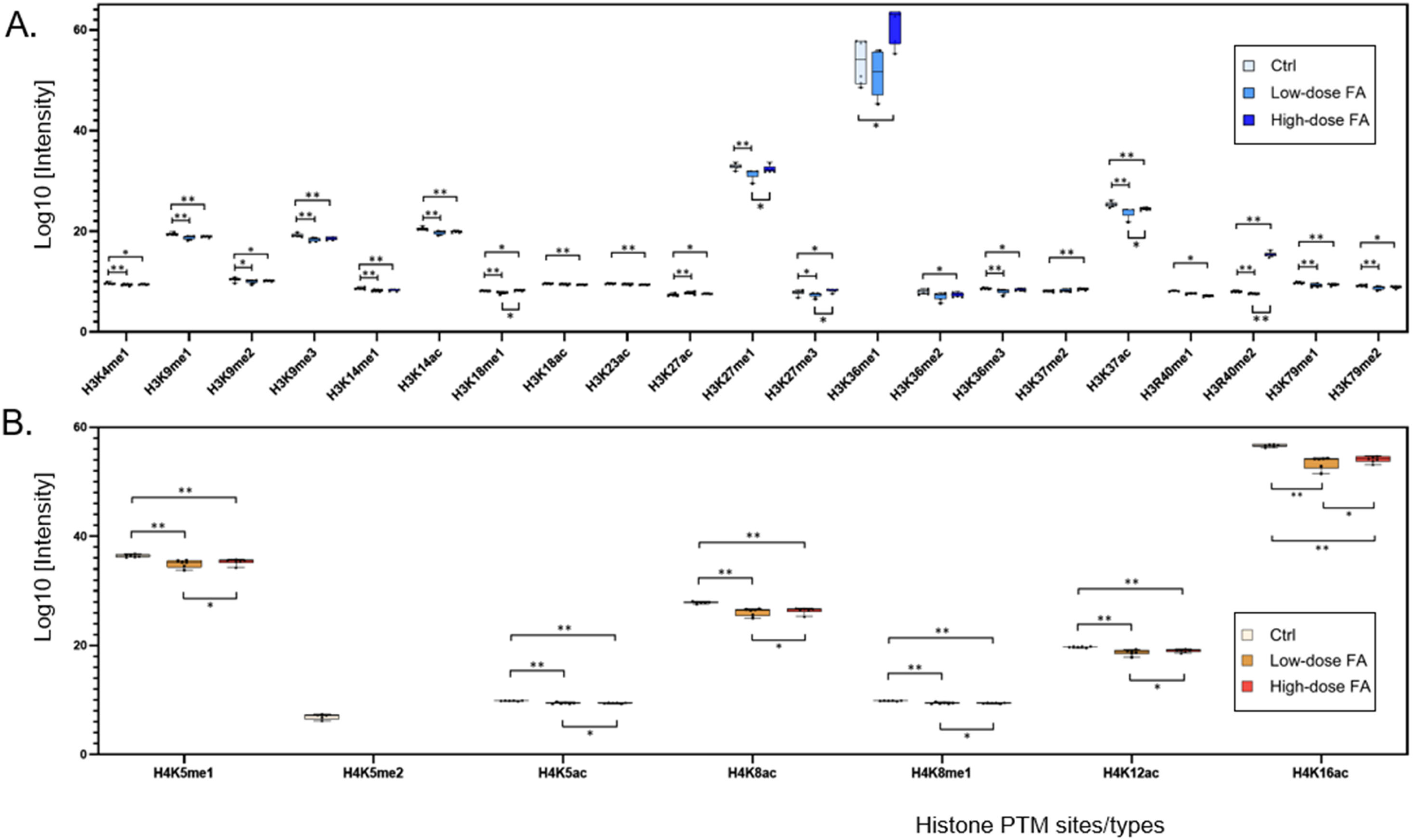
Comparison of PTM site/type mapping on histones H3 (A) and H4 (B) across control, low-dose, and high-dose FA exposure groups (N = 6). Statistical significance is indicated as \**p* < 0.05 and \*\**p* < 0.01.

## 4. Discussion

In this study, we investigated the impact of FA on histone post-translational modifications (PTMs), focusing on histone H3 and H4 methylation and acetylation. Using high-resolution liquid chromatography-tandem mass spectrometry (LC-MS/MS)-based proteomics, we provided a detailed and quantitative analysis of these modifications. Notably, our findings highlighted the sensitivity of histone PTMs to FA exposure, revealing distinct patterns of alteration in both methylation and acetylation levels. This work also facilitated the understanding of FA-induced epigenetic changes by employing intensity-based peptide-level analysis, offering a novel perspective on the epigenetic landscapes affected by FA exposure. These results underscore the potential of FA to disrupt chromatin dynamics and gene expression regulation, shedding light on exploring its mechanisms in carcinogenesis. The systemic alteration of histone methylation and acetylation on H3 and H4 might be the key linkages between FA exposure and multiple cellular processes.

### 4.1. The effects of FA on H3 methylation

Peptide analysis in this study directly revealed impacts of FA on methylation states of H3K4 and H3K79, which was displayed consistently as suppression, regardless of low-dose or high-dose FA exposure (Figure 3A; 3B; 3C). Methylation of H3K79 is involved in the regulation of telomeric silencing, cellular development, cell-cycle checkpoint, DNA repair, and regulation of transcription (Farooq et al., 2016). H3K79 is methylated by methyltransferase DOT1L in a non-processive manner (Frederiks et al., 2008; Min et al., 2003) using S-adenosylmethionine (SAM) as a cofactor. FA exposure causes SAM deficiency (Pham et al., 2023), which indicated that FA exposure is likely to reduce DOT1L activity due to SAM deficiency and consequently inhibit H3K79 methylation. H3K79me2 is the epigenetic signature for mixed lineage leukemia, as its abnormal amount and distribution were associated with MLL-AF9 target loci (Bernt et al., 2011). DOT1L has been implicated in the development of leukemias bearing translocations of the Mixed Lineage Leukemia gene (Bernt et al., 2011), suggesting that FA-induced decrease in H3K79me2 might be a critical contributor to leukemogenesis.

H3K4me1 is a well-established mark of both poised and active enhancers (Heintzman et al., 2007). A recent study demonstrates that H3K4me1 enhances enhancer-promoter contacts and enhancer-driven transcription during cell differentiation (Kubo et al., 2024). MLL2, a major mammalian histone methyltransferase that mono-methylates H3K4, is a tumor suppression gene in non-Hodgkin lymphoma (Ortega-Molina et al., 2015; J. Zhang et al., 2015). MLL2, also known as histone lysine N-methyltransferase (KMT2D), is frequently mutated in follicular lymphoma (FL) and diffuse large B cell lymphoma (DLBCL) (Ortega-Molina et al., 2015; J. Zhang et al., 2015). The loss of MLL2 promoted lymphoma development by impairing H3K4 methylation and altering gene expression in pathways critical for B cell activation and differentiation. Reduction of H3K4me1 by FA exposure (Figure 3A and 8A) suggests a potential underlying mechanism for lymphoma induced by FA.

Methylation of H3K9 has been implicated in heterochromatin formation and gene silencing (Bannister et al., 2001). H3K9me2 and H3K9me3 specifically bind chromodomain proteins, such as the heterochromatin protein 1 family, to form a higher-order architecture of heterochromatin, leading to gene repression (Bannister & Kouzarides, 2011; Xhemalce et al., 2011). Lower levels of H3K9me1 were also observed in less active promoters around the transcription start sites (Barski et al., 2007). FA exposure induced a significant decrease in global H3K9me1, H3K9me2, and H3K9me3 (Figure 4A and Figure 8A). The level of H3K27me3, a well-known marker of transcription repression (M.-R. Pan et al., 2018; Shen et al., 2008), decreased at low FA doses but increased at high doses. These results suggest that FA might contribute to cancer development through disrupting heterochromatin structure and dysregulation of cancer-related genes. In fact, numerous studies have reported an association between H3K27Me3 and various cancers. Notably, the expression of H3K27Me3 is elevated in NPC relative to normal nasopharyngeal epithelial tissues, which contributes to enhanced metastatic potential (Cai et al., 2011), which proved the histone modification might be another potential mechanism of FA-induced NPC. Moreover, increased H3K27me3 level was confirmed to correspond to the progression and proliferation rate of both primary and immortalized acute myeloid leukemia cells (Li et al., 2018).

FA exposure also reduced the global levels of H3K36 methylation, in particular H3K36me3, one of the extensively studied histone marks. H3K36me3 has been reported to function in the regulation of transcriptional activity, transcription elongation, and alternative splicing. Additionally, H3K36me3 also contributes to DNA replication, recombination, and repair (Sun et al., 2020; Wagner & Carpenter, 2012). Loss of H3K36me3 induced multiple human cancers, including clear cell renal cell carcinoma (ccRCC), high-grade gliomas, and hematopoietic malignancies (Fahey & Davis, 2017; Xie et al., 2022). FA exposure is associated with an increased risk of leukemia. Interestingly, in acute leukemia, SETD2 mutations, a histone H3K36 methyltransferase, cause a global loss of H3K36me3, facilitating leukemia stem cell self-renewal and promoting both initiation and progression of the malignancy. In chronic myelogenous leukemia (CML), H3K36me3 is identified as a key histone modification regulating gene expression changes associated with leukemogenesis. These findings establish SETD2 mutation and loss of H3K36me3 as critical components in leukemia pathogenesis, providing potential mechanisms underlying the increased risk of leukemia associated with FA exposure.

### 4.2. The effects of FA on H3 acetylation

FA exposure lowered the acetylation of H3K14, K18, K23, and K37, while it increased the level of H3K27ac. H3K27ac is the only modification upregulated by FA exposure at both low- and high-dose. Whereas it is unknown how these alterations affect the expression of individual genes during pathogenesis, changes in global histone acetylation are detected in different types of cancers and linked to cancer development and progression. For example, a reduced level of H3K18Ac has been observed in the prostate, kidney, breast, lung, and pancreatic cancers (Elsheikh et al., 2009; Manuyakorn et al., 2010; Seligson et al., 2005, 2009). Furthermore, decreased H3K18Ac levels have been identified in low-grade prostate cancer patients, particularly those with poor prognoses and a high likelihood of tumor recurrence (Seligson et al., 2005). H3K18Ac is considered a marker of cancer progression as deacetylation of H3K18Ac is crucial for maintaining essential features of cancer cells (Barber et al., 2012).

H3K27ac is a well-known marker of active enhancers and plays a role in enhancer RNA transcription (Kang et al., 2021). Increased H3K27ac level was detected in epithelial ovarian cancer, which was associated with decreased overall survival and progression-free survival (Bauer et al., 2021). Moreover, H3K27ac has been shown to enhance cancer cell survival and proliferation in various cell lines, including esophageal squamous cell carcinoma, lung squamous cell carcinoma, and hepatocellular carcinoma cell lines (Miziak et al., 2024). Notably, it was demonstrated that H3K27ac may play a critical role in acute myeloid leukemia (AML), by activating enhancers that maintain leukemic stem cell (LSC) oncogenic potential. Inhibition of EP300, a histone acetyltransferase, disrupted H3K27ac at enhancers and impaired LSC maintenance, highlighting the potential of targeting EP300 and H3K27ac as a therapeutic strategy in AML (F. Pan et al., 2023). These results suggest that FA exposure may contribute to AML development by increasing the level of H3K27ac.

### 4.3. The effects of FA on H4 acetylation

Our results showed that FA exposure reduced the levels of H4K5ac, H4K8ac, H4K12ac, and H4K16ac at both low- and high-dose. We have previously demonstrated that FA exposure inhibits H4K12ac probably through the formation of FA-lysine adduct at the sites, preventing the sites from being physiologically acetylated. The reduction of H4K12ac, alongside with downregulation of acetylation of N-tail H3, contributed to FA-induced inhibition of chromatin assembly (Chen et al., 2017). H4K5ac also plays an important role in histone translocation and nucleosome assembly. Thus, the results from this study further resonate that FA exposure may compromise chromatin assembly by reducing the levels of H4K5ac, H412ac and H3 N-terminal tail acetylation. All histone acetylations are also involved in transcriptional activation. Therefore, reduced levels of H4K5ac, H4K8ac, H4K12ac, and H4K16ac may contribute to FA toxicity by inducing aberrant transcription. Among these H4 acetylations, the loss of H4K16ac is considered a common hallmark of human cancers (Fraga et al., 2005), since the loss predominantly occurred at the H4K16, but not at H4K5, H4K8, or H4K12, in 36 primary tumors and 25 cancer cell lines including 4 leukemia cell lines. Therefore, reduction of H4K16ac may also contribute to FA-induced hematopoietic malignancy.

### 4.4. Implications in neurodegenerative diseases

FA-induced histone modifications may constitute a critical mechanism underlying the pathogenesis of neurodegenerative diseases, as the perturbed histone modification landscapes identified in this study closely parallel epigenetic signatures observed in patients and rodent models of neurodegenerative diseases. In the context of Alzheimer’s disease (AD), a pronounced depletion of global H3K18ac and H3K23ac levels has been reported in the temporal lobes of affected individuals (Zhang et al., 2012). Substantial reductions in H3K9me2 and H4K12ac levels were detected in the hippocampal cornu ammonis 1 (CA1) region in a cohort of 47 Alzheimer’s disease (AD) cases (Hernández-Ortega et al., 2016). This finding is consistent with a report indicating that the level of histone H4 acetylation is reduced by 50% in APP/PS1 mice compared to their wild-type littermates (Francis et al., 2009). Interestingly, the restoration of H4 acetylation through the use of HDAC inhibitors was shown to improve learning in mice, suggesting a potential role for the loss of H4Kac in cognitive impairment. Epigenomic landscapes and their correlation with transcriptional changes during neurodegeneration in the hippocampus of the CK-p25 mouse model of AD and CK littermate controls revealed notable differences in H3K27ac levels, which included 2456 increased-level peaks and 2154 decreased-level peaks (Gjoneska et al., 2015). Furthermore, AD-related genetic variants identified through GWAS were enriched in orthologues of increased-level enhancers marked by H3K27ac. A genome-wide analysis of H4K16ac, a histone modification associated with aging, demonstrated that normal aging is characterized by increased H4K16ac levels, whereas a significant loss of H4K16ac was observed near genes associated with aging and AD in the lateral temporal lobes of AD individuals compared to younger, cognitively normal controls (Nativio et al., 2018). Additionally, this same study identified a strong association between AD-related GWAS single-nucleotide polymorphisms and the expression quantitative trait loci within regions exhibiting significant H4K16ac alterations (Nativio et al., 2018). The aforementioned histone marks related to AD pathogenesis were found matched to the FA-induced histone patterns (Figure 8A and 8B), indicating that correlations between FA exposure and AD could be established through histone-related epigenetic mechanisms.

## 5. Conclusions

FA exposure elicits extensive and systemic perturbations in histone methylation and acetylation, specifically targeting histones H3 and H4. Through the application of cutting-edge high-resolution liquid chromatography-tandem mass spectrometry-based proteomics, peptide sequence analysis revealed that FA exposure induces hypomethylation at key residues, including H3K4 and H3K79, while significantly disrupting the methylation and acetylation landscape at H3K9, H3K14, H3K18, H3K23, H3K27, H3K36, and H4K37. Notably, mono-methylation and di-methylation of arginine residues on histone H3 were also markedly affected. On histone H4, profound alterations were observed in the acetylation states of lysine residues K5, K8, K12, and K16. Intricate mapping of single PTM types and sites further revealed that the epigenetic modifications induced by FA closely mimic histone modification patterns previously implicated in cancers, including nasopharyngeal cancer and leukemia, as well as AD, underscoring changes in histone modification as a potential epigenetic mechanism driving FA-associated pathological outcomes.

## Acknowledgements

This work was supported by the National Institute of Environmental Health Sciences (NIEHS) of the National Institutes of Health (NIH) under grant number 1R01ES033160.

